# Cortical Responses Time-Locked to Continuous Speech in the High-Gamma Band Depend on Selective Attention

**DOI:** 10.1101/2023.07.20.549567

**Authors:** Vrishab Commuri, Joshua P. Kulasingham, Jonathan Z. Simon

## Abstract

Auditory cortical responses to speech obtained by magnetoencephalography (MEG) show robust speech tracking to the speaker’s fundamental frequency in the high-gamma band (70-200 Hz), but little is currently known about whether such responses depend on the focus of selective attention. In this study 22 human subjects listened to concurrent, fixed-rate, speech from male and female speakers, and were asked to selectively attend to one speaker at a time, while their neural responses were recorded with MEG. The male speaker’s pitch range coincided with the lower range of the high-gamma band, whereas the female speaker’s higher pitch range had much less overlap, and only at the upper end of the high-gamma band. Neural responses were analyzed using the temporal response function (TRF) framework. As expected, the responses demonstrate robust speech tracking of the fundamental frequency in the high-gamma band, but only to the male’s speech, with a peak latency of approximately 40 ms. Critically, the response magnitude depends on selective attention: the response to the male speech is significantly greater when male speech is attended than when it is not attended, under acoustically identical conditions. This is a clear demonstration that even very early cortical auditory responses are influenced by top-down, cognitive, neural processing mechanisms.

## 1 INTRODUCTION

Time-locked auditory responses are one mechanism by which the auditory system preserves temporal information about sounds. For example, subcortical responses to voiced sections of speech time-lock to the speaker’s fundamental frequency (F0), whether ;2100 Hz for a typical male voice (Skoe and Kraus, 2010) or ;2200 Hz for a typical female voice (Lehmann and Schönwiesner, 2014), and have been measured via the frequency following response (FFR)(Kraus et al., 2017). As neural responses propagate up the auditory pathway, characteristic time-locking frequencies are generally observed to decrease. For example, cortical responses time-lock to the envelope of the speech most strongly below *∼* 10 Hz (Ahissar et al., 2001; Luo and Poeppel, 2007). Nevertheless, recent FFR studies have observed cortical time-locked responses at rates often associated with subcortical processing, ≳100 Hz, using responses measured from magnetoencephalography (MEG) (Coffey et al., 2016; Gorina-Careta et al., 2021), and electroencephalography (EEG) (Bidelman, 2018). However, even the highest frequencies associated with cortical phase locking are substantially lower than those seen from subcortical sources (typically with EEG).

The FFR obtained from the average of many (e.g., thousands of) responses to a repeated auditory stimulus has been used to provide insight into the representation of speech in the auditory periphery and the fidelity of sound encoding in the brain (Basu et al., 2010; Kraus et al., 2017). Modulations of the FFR strength and consistency can be used to study cognitive processes such as learning (Skoe et al., 2013), selective attention (Lehmann and Schönwiesner, 2014; Holmes et al., 2017), level of attention (Price and Bidelman, 2021), intermodal (auditory vs. visual) attention (Hartmann and Weisz, 2019), and the effect of familiar vs. unfamiliar background language (Presacco et al., 2016; Zan et al., 2019). These studies demonstrate that FFRs can be affected by top-down auditory processes, though it is not clear how much of the FFR modulation is due to subcortical vs. cortical sources (Gnanateja et al., 2021; Gorina-Careta et al., 2021).

The FFR, in order to be averaged over so many trials, uses many repetitions of a short stimulus (e.g., a single speech syllable). In contrast, temporal response functions (TRFs), used here, characterize neuronal responses to speech using single long-duration trials of continuous speech (Lalor et al., 2009; Ding and Simon, 2012). While TRF analysis is most often applied to low frequency cortical responses (Brodbeck and Simon, 2020), TRFs obtained with MEG have recently also been used to investigate cortical responses to speech in the high-gamma range (70-200 Hz)(Kulasingham et al., 2020; Schüller et al., 2023a), i.e., for frequencies similar to those investigated using cortical FFR, showing a single response peak with latency *∼* 40 ms, indicating a focal neural origin in primary auditory cortex (see also Kegler et al. (2022) for EEG). The present study extends the work of Kulasingham et al. (2020) by applying high-gamma TRF analysis of MEG responses to subjects listening to speech from male and female speakers in single-speaker and “cocktail-party” (competing speaker) paradigms.

The present study also uses single-speaker conditions to allow comparison of subjects’ responses to both male (F0 ≳ 100 Hz) and female speech (F0 ≳ 200 Hz) in isolation. Prior work has posited that high-gamma cortical responses may reflect the processing of F0 and related features in a speech stimulus (Guo et al., 2021). Additionally, Kulasingham et al. (2020) found that high-gamma cortical responses were driven mainly by the segments of speech with F0 » 100 Hz and that responses to F0 above 100 Hz were not easily detected. This suggests that responses to speech from a typical female speaker (average F0 100 Hz) may be reduced in comparison to responses to a male speaker (average F0 *∼* 100 Hz). Moreover, many recent studies on high-gamma cortical responses to speech only use stimuli from male speakers in their experimental design (Kulasingham et al., 2020; Canneyt et al., 2021a; Gnanateja et al., 2021; Guo et al., 2021; Kegler et al., 2022; Schüller et al., 2023b). This may be because typical male speakers have a lower F0 than typical female speakers, and stronger responses are evoked by speech with a lower F0. Indeed, Canneyt et al. (2021b) investigated responses to stimuli from both male and female speakers and observed that high-gamma cortical response strength was inversely related to F0.

The competing speakers conditions used here allow the investigation of how these fast cortical responses change depending on top-down influences such as task specificity and selective attention. The use of both a male and female speaker removes much of the ambiguity as to the source of the responses due to the considerable gap between the speakers’ fundamental frequency bands with the aim of enhancing responses to the male speech stream which can be assessed for attentional effects. In humans it is seen widely that auditory low frequency (;S10 Hz) time-locked cortical responses depend on selective attention, whether for simple sounds (Hillyard et al., 1973; Elhilali et al., 2009; Holmes et al., 2017) or speech (Lalor et al., 2009; Ding and Simon, 2012). To what extent selective attention changes response properties in early latency primary auditory cortex, as opposed to secondary auditory areas and beyond, is not yet well understood. Using invasive intracranial EEG (iEEG) recordings, effects of selective attention have been observed for simple stimuli (Bidet-Caulet et al., 2007) but not for competing speakers (O’Sullivan et al., 2019). From MEG studies there is recent evidence for selective attention affecting the low frequency response properties of very early auditory cortex during a competing speaker task (Brodbeck et al., 2020), but the effect is small and occurs only under limited conditions.

Thus, the main focus of the present study concerns two primary research questions. Firstly, what differences are there in high-gamma cortical responses between the cases of male (F0 ≳ 100 Hz) vs female (F0 ;2 200 Hz) speech? Secondly, do early (*∼* 40 ms latency) high-gamma cortical responses to speech, putatively arising only from primary auditory cortex (Simon et al., 2022), depend on selective attention? Both these questions are addressed by analyzing MEG recordings of subjects listening to single male and female voices, and to the same voices presented simultaneously but with the task of selectively attending to only one or the other.

## 2 MATERIALS AND METHODS

### 2.1 Data

The data set analyzed here was previously obtained and analyzed in an earlier study that investigated differing cortical responses between spoken language and arithmetic using two different speakers (Kulasingham et al., 2021). The data are available at https://doi.org/10.13016/xd2i-vyke and the code is available at https://github.com/vrishabcommuri/mathlang-highgamma.

### 2.2 Participants

The data set comprises MEG responses recorded from 22 individuals (average age 22.6 years, 10 female, 21 right handed) who were native English speakers. Individuals underwent a screening in which they self-reported any known hearing issues, and a brief MEG pre-experiment recording to verify that auditory cortical responses to 1 kHz tone pips were present and normal. No subjects were excluded on either ground. The participants provided written informed consent and received monetary compensation. The experimental procedure was approved by the Internal Review Board of the University of Maryland, College Park.

### 2.3 Data Acquisition and Preprocessing

The data were collected from subjects using a 157 axial gradiometer whole head KIT (Kanazawa Institute of Technology) MEG system with subjects resting in the supine position in a magnetically shielded room (Vacuumschmelze GmbH & Co. KG, Hanau, Germany). The data were recorded at a sampling rate of 1 kHz with an online 200 Hz low pass filter with a wide transition band above 200 Hz and a 60 Hz notch filter. Data were preprocessed in MATLAB by first automatically excluding saturating channels and then applying time-shift principal component analysis (TSPCA) (de Cheveigné and Simon, 2007) to remove external noise, and sensor noise suppression (SNS) (de Cheveigné and Simon, 2008) to suppress channel artifacts. Two of the sensor channels were excluded during the preprocessing stage.

The denoised MEG data were filtered from 70 to 200 Hz using an FIR bandpass filter with 5 Hz transition bands and were subsequently downsampled to 500 Hz. Independent component analysis (ICA) was then applied to remove artifacts such as heartbeats, head movements, and eye blinks.

The subsequent analyses were performed in Python using the mne-python (1.3.1) (Gramfort et al., 2013) and eelbrain (0.38.4) (Brodbeck et al., 2019) libraries, and in R using the lme4 (1.1-21) (Bates et al., 2015) and buildmer (2.8) (Voeten, 2023) packages.

### 2.4 Neural Source Localization

Prior to the data collection, the head shape of each subject was digitized using a Polhemus 3SPACE FASTRAK system, and subject head position in the MEG scanner was measured before and after the experiment using five marker coils. The marker coil locations and the digitized head shape were used to co-register the template FreeSurfer “fsaverage” brain (Fischl, 2012) using rotation, translation, and uniform scaling.

Source localization was performed using the mne-python software package. First, a volume source space was composed from a grid of 7-mm sized voxels. Then, an inverse operator was computed, mapping the sensor space to the source space using minimum norm estimation (MNE) (Hämäläinen and Ilmoniemi, 1994) and dynamic statistical parametric mapping (dSPM) Dale et al. (2000) with a depth weighting parameter of 0.8 and a noise covariance matrix estimated from empty room data. The result of the localization procedure was a single 3-dimensional current dipole centered within each voxel.

The Freesurfer ‘aparc+aseg’ parcellation was used to define a cortical region of interest (ROI). The ROI consisted of voxels in the gray and white matter of the brain that were closest to the temporal lobe – Freesurfer ‘aparc’ parcellations with labels ‘transversetemporal’, ‘superiortemporal’, ‘inferiortemporal’, and ‘bankssts’. All analyses were constrained to this ROI to conserve computational resources.

### 2.5 Stimuli

Subjects listened to isochronous (fixed-rate) speech from two synthesized voices – one male and one female. Speech was generated using the ReadSpeaker synthesizer with the “James” and “Kate” voices (https://www.readspeaker.com). Two kinds of speech stimuli were created: ‘language’ stimuli that consisted of four-word sentences, and ‘arithmetic’ stimuli that consisted of five-word equations. The word rate of the arithmetic stimuli was faster than that of the sentence stimuli so that neural responses to each could be separated in the frequency domain and so that each stimulus was 18 seconds in duration. The stimulus files are available in the same repository as the data: https://doi.org/10.13016/xd2i-vyke.

### 2.6 Experimental Design

The experiment was divided into two conditions. In the first condition (‘single-speaker’), subjects listened to speech from either the male or the female speaker; and in the second condition (‘cocktail-party’), subjects listened to both speakers concurrently and were instructed to attend to only one. Each condition was conducted in blocks: four single speaker blocks (2 *×* 2: male and female, sentences and equations) followed by eight cocktail party blocks. At the start of each cocktail party block, the subject was instructed as to which stimulus to attend to, and was asked to press a button at the end of each trial to indicate whether a deviant was detected. The subjects were generally able to attend to the instructed speaker (Kulasingham et al., 2020). The order in which blocks were presented was counterbalanced across subjects.

### 2.7 Stimulus Representations

In accordance with the methods of Kulasingham et al. (2020), two predictors (i.e., stimulus representations) were used, one capturing the high-frequency envelope modulations and another capturing the stimulus carrier (also called temporal fine structure; TFS). The broad rationale for using these two predictors is to allow comparisons with the analogous varieties of FFR: FFR_ENV_ and FFR_TFS_ (Coffey et al., 2019). Figure 1 illustrates the procedure for extracting both predictors from a stimulus waveform.

**Figure 1.**
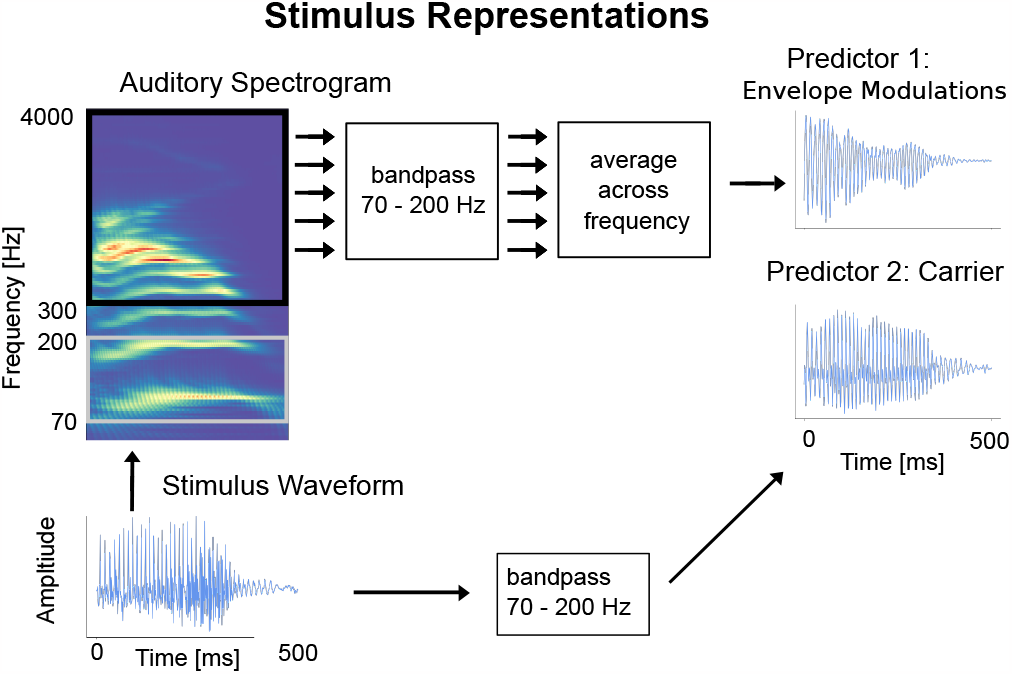
Illustration of how the carrier and envelope modulations predictors are extracted from an auditory stimulus. The raw stimulus waveform is shown in the bottom-left corner. **Envelope modulations predictor:** to generate the envelope modulations predictor, starting with the raw waveform and following the arrows up and to the right, first an auditory spectrogram is generated using a model of the auditory periphery (Yang et al., 1992). Then, the acoustic envelope in each frequency bin in the range 300 to 4000 Hz is bandpassed in the high-gamma range (70-200 Hz), and the average is then computed across the channels. The result is a single time-series signal. **Carrier predictor:** to generate the carrier predictor, following the arrows to the right, the raw stimulus waveform is simply bandpass filtered to the high-gamma range. The result is a second single time-series signal. (Figure reproduced with permission from Kulasingham et al. (2020).)

#### 2.7.1 Carrier Predictor

The carrier predictor is a representation of the speech signal components within the high-gamma band. In particular, the fundamental frequency of voiced speech is directly encoded by this representation. The inclusion of this stimulus representation as a predictor in our model enables us to examine how much of the neural response is a consequence of cortical entrainment to the high-gamma frequencies of the stimulus waveform itself.

To create the carrier predictor, each stimulus was first resampled to a frequency of 500 Hz to reduce the ensuing computation required. Prior to downsampling, an anti-aliasing FIR prefilter with 500 Hz cutoff and 5 Hz transition band was applied to the data. This resampled signal was then bandpass filtered in the high-gamma range of 70-200 Hz using an FIR filter with 5 Hz transition band. Finally, the signal was standardized (i.e., mean subtracted and normalized by the standard deviation) to produce the carrier predictor. Standardized carrier predictors for each stimulus in the condition were concatenated to form one long-form carrier predictor per condition.

#### 2.7.2 Envelope Modulations Predictor

In contrast to the carrier predictor, which extracts high-gamma band components directly from the stimulus, the envelope modulations representation captures high-gamma band modulations of higher frequency bands present in the stimulus. Higher frequency bands capture harmonic content that cannot be derived from the fundamental frequency alone, but which, for voiced sections of speech, is modulated at the rate of the fundamental frequency of the voicing due to the inherent non-linearities of the auditory system. We include the envelope modulations predictor in our model to assess cortical entrainment to the high-gamma band envelope modulations of these higher frequency signals.

To create the envelope modulations predictor, the speech was transformed into an auditory spectrogram representation at millisecond time resolution using a model of the auditory periphery (Yang et al., 1992) (http://nsl.isr.umd.edu/downloads.html). The model uses a bank of 128 overlapping *Q*_10dB_ *≈* 3 bandpass filters uniformly distributed along a logarithmic frequency axis over 5.3 oct (24 filters/octave); other details of the model, including the hair-cell stage and lateral inhibition with half-wave rectification are described in Chi et al. (2005).

The auditory spectrogram produced by the model is a two-dimensional matrix representation of the acoustic envelope over time for different frequency bins. The spectrogram frequency bins in the range 300-4000 Hz were selected, resulting in a time-series for each frequency bin: the time course of the acoustic power in the signal in that band. The range 300-4000 Hz was chosen in order to effect a clear separation between the lowest frequency in the predictor the upper end of the high-gamma range (200 Hz) and because the stimulus was presented through air tubes which attenuate frequencies above 4000 Hz (Kulasingham et al., 2021). Each time-series was filtered to the high-gamma range in the same method as the carrier predictor, using an FIR filter with a 70-200 Hz passband. The time-series signals were then averaged across frequency bins, and the resulting signal standardized, to produce a single time-series – the envelope modulations predictor.

### 2.8 TRF Estimation

The simplest model of a single temporal response function (TRF) is given by

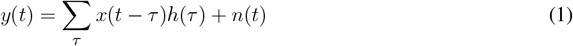

where *x*(*t − τ*) is the time-shifted predictor signal (e.g., high-frequency envelope modulations or carrier) at time lag *τ* ; *h*(*τ*) is the TRF at time lag *τ* ; *y*(*t*) is the MEG measured response signal; and *n*(*t*) is the residual noise (i.e., everything not captured by convolving the predictor and TRF).

From equation 1, we see that the TRF *h* is simply the impulse response of the neural system with predictor input *x* and with MEG measured response output *y*. The TRF can be interpreted as the average time-locked neural response to continuous stimuli (Lalor and Foxe, 2010).

#### 2.8.1 Single-Speaker Model

In the present study, a more complex model with two predictors was used for the single-speaker condition

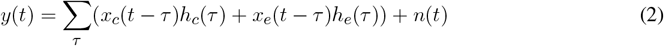

where *x*_*c*_ and *x*_*e*_ are, respectively, the carrier and envelope modulations predictors derived from the single-speaker stimulus, and *h*_*c*_ and *h*_*e*_ are the corresponding TRFs.

#### 2.8.2 Cocktail-Party Model

A TRF model with four predictors – the carrier and envelope modulations for the attended speaker and the carrier and envelope modulations for the unattended speaker – was used for the cocktail-party conditions.

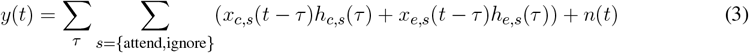

where predictors and TRFs are similar to the single-speaker model in 2, with the additional subscript *s* indicating the attended and unattended speaker. TRFs corresponding to the male and female speakers were analyzed separately, but TRFs corresponding to ‘attend language’ and ‘attend arithmetic’ were pooled together within each speaker.

#### 2.8.3 Estimation Procedure

The parameters for each TRF model were estimated jointly such that the ordering of the predictors did not affect the estimates, enabling predictors to compete to explain the variance in the data. Predictors that contributed more to the neural response had larger TRFs. TRFs were estimated for time lags from -40 to 210 ms using boosting with cross-validation via the ‘boosting’ routine from the eelbrain library (Brodbeck et al., 2019). Overlapping bases of 4 ms Hamming windows with 1 ms spacing were employed to promote smoothly varying responses.

Since the source space MEG responses are 3-dimensional current vectors, the estimated TRFs also comprise vectors that span 3 spatial dimensions. The L2 norm (amplitude) of each vector in the TRF was taken at each time instance, resulting in a 1-dimensional time-series for each TRF—one TRF per source space voxel—thereby simplifying the interpretation and visualization of the results.

### 2.9 F0 Analysis

To investigate the extent to which the MEG responses in our study were affected by speaker F0, a simple comparison was conducted whereby the time-averaged F0 of each speaker was extracted using Praat (Boersma and Weenink, 2022) and then compared to the amplitude of the TRFs.

### 2.10 Statistical Tests

To determine whether peaks in the estimated TRFs were induced by time-locked neural responses to the predictors and not simply obtained by chance, a null model was created by circularly time-shifting the predictors and recomputing TRFs using the shifted predictors. This procedure enables us to disentangle responses to the typical temporal structure of the predictor from responses that time-lock to the predictor. Three shifted versions of each predictor were produced by shifting in increments of one-fourth of the total duration of the original predictor, resulting in three null-model TRFs for each original TRF. Cluster-based permutation tests (Nichols and Holmes, 2001) with Threshold Free Cluster Enhancement (TFCE) (Smith and Nichols, 2009) were used to test for significance across the TRF peak regions over the average of the three null models and to account for multiple comparisons. Significance for all tests was set at the 0.05 level.

To test that the TRFs were better than chance at predicting the MEG responses, we compared the prediction accuracy of the TRF model to the average prediction accuracy of the three null models. Since all predictors were fit jointly, this results in one prediction accuracy per voxel per model. Because each subject was mapped individually to the ‘fsaverage’ brain, individual variation was mitigated by smoothing the voxel prediction accuracies over the source space using a Gaussian window with 5 mm standard deviation. Cluster-based permutation tests with TFCE were used to test for significance across the cortical region of interest.

TRFs were computed for each source voxel as a time-varying, three-dimensional current dipole that varies over time lags. For each TRF vector, its amplitude was compared to the average of three null models across subjects at each time lag. Time lags for which the true model amplitude was significantly greater than the average null model were determined using a one-tailed test with paired sample *t*-values and TFCE.

To assess differences in TRF peak amplitude across conditions (single-speaker and cocktail-party) two linear mixed effects models were used. Prior to fitting, the average of the three null models was subtracted from each TRF; this had the effect of subtracting off the noise floor of each TRF, thereby facilitating a more direct comparison of peak amplitudes. From the result, the peak amplitudes in the range 20-50 ms were extracted. For each condition, two models were developed: one maximal model that attempts to account for as many fixed (population-level) and random (subject-level) effects as possible in the data, and a reduced model that was obtained by pruning effects from the maximal model that failed to significantly explain variance in the data. Maximal models are the largest possible models that will still converge and were obtained using the R package buildmer. Reduced models were then obtained for each condition using buildmer’s backward elimination protocol.

The linear mixed effects models were fit to the TRF peak amplitudes and incorporated the following categorical inputs: predictor type (either carrier or envelope modulations) and speaker gender (either male or female) in the single-speaker model; and predictor type (either carrier or envelope modulations) and attention focus (either attend or ignore male speaker) in the cocktail-party model. The target maximal model for buildmer was in both cases obtained by setting all crossed terms as fixed and random effects:

*SS*_maximal_ : peak amplitude *∼* predictor type*×*speaker gender+(predictor type *×*speaker gender| subject)

*CP*_maximal_ : peak amplitude *∼* predictor type*×*attention focus+(predictor type *×*attention focus| subject)

## 3 RESULTS

### 3.1 F0 Analysis

The average F0 for each speaker was computed for voiced regions of speech over all trials:

- Male speaker: average F0 of 95 Hz (std. dev. 8 Hz)
- Female speaker: average F0 of 168 Hz (std. dev. 10 Hz)

Kulasingham et al. (2020) found that neural responses are diminished for F0 above 100 Hz. Because the male speaker’s average F0 is below 100 Hz, and the female speaker’s average is well above 100 Hz, we anticipate stronger high-gamma cortical responses for the male speaker than the female speaker.

### 3.2 TRF Response Estimation

To validate the extent to which the estimated TRFs can predict the neural responses from the predictor signals, a prediction accuracy is computed for each TRF. The prediction accuracy is the correlation coefficient between the normalized predicted and true neural responses for each TRF. Since a TRF was estimated for each voxel in the source space, this assesses which cortical regions were best predicted by the TRF model.

Prediction accuracies were computed for the single-speaker and cocktail-party models (single-speaker: mean = 0.0149, std = 0.0065; cocktail-party: mean = 0.0132, std = 0.0061). The prediction accuracies for the average of the three null models (single-speaker null: mean = 0.0123, std = 0.0057; cocktail-party null: mean = 0.0115, std = 0.0059) were compared to the original models by means of a one-tailed test with paired sample *t*-values and TFCE for each of the two conditions. A large portion of the voxels showed a significant increase in prediction accuracy over the null model (single-speaker: *t*_max_ = 7.372, *p <* 0.001, cocktail-party: *t*_max_ = 5.055, *p <* 0.001; see Figure 2).

**Figure 2.**
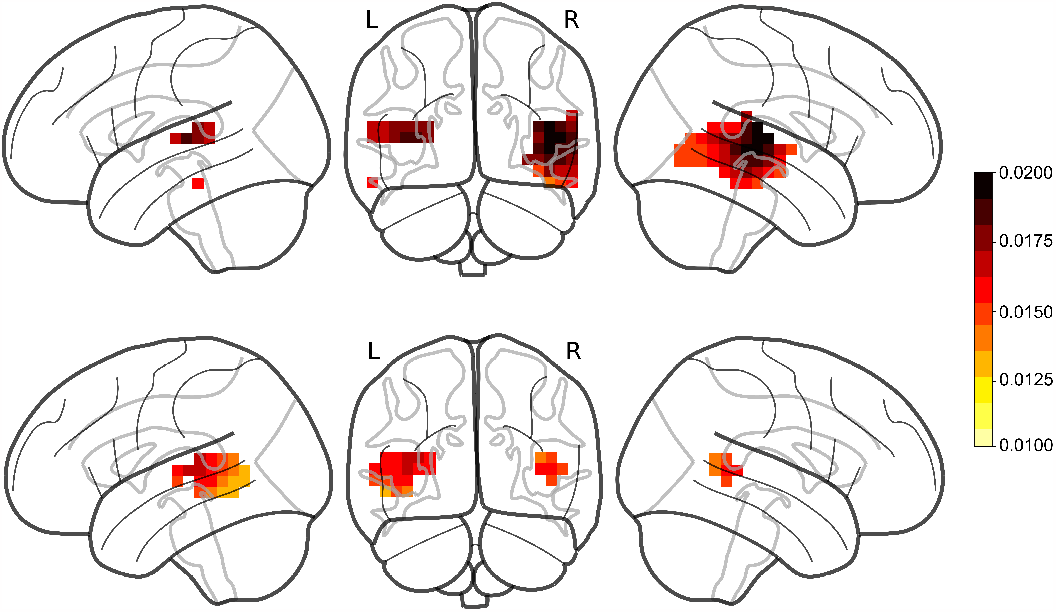
Prediction accuracies for male single-speaker (**Top**) and cocktail-party (**Bottom**) models. Red regions denote voxels where the TRF model produced a prediction accuracy that was significantly greater than that of the noise within the ROI. TRFs to female speech (not shown) did not produce significant responses in any voxels.

No significant voxels were identified in TRFs for the female single speaker (single-speaker: *t*_max_ = 3.436,*p* = 0.055).

### 3.3 Single-Speaker TRFs

Figure 3 shows the various TRFs, averaged across voxels and subjects, and latency ranges for which the TRFs were significantly greater than the noise floor. In total, four TRFs were computed for the single-speaker scenario: carrier and envelope TRFs for male and female speakers.

**Figure 3.**
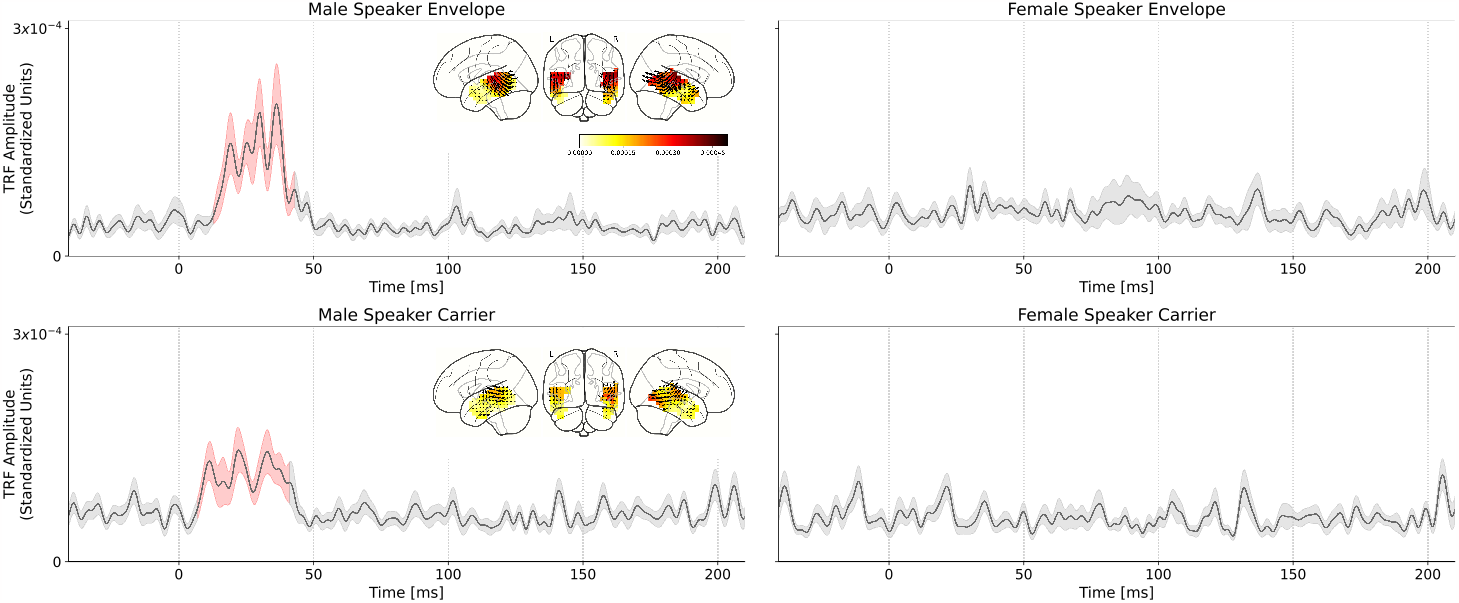
Comparison of male speech and female speech TRFs for the single speaker conditions. Solid black lines indicate the TRF grand average (over TRF amplitude, averaged across voxels in the ROI); shaded regions indicate values within one standard error of the mean. Red shading indicates TRF values significantly above the noise floor. The distribution of TRF vectors in the brain at the time with the maximum significant response is plotted as an inset for each TRF. **Top Left:** average TRF of the envelope modulations predictor derived from the male speaker stimulus. Note the large significant response at ~30-50 ms in the TRF which indicates a consistent, time-locked neural response to the speech envelope modulations at a *∼* 30-50 ms latency. **Top Right:** average TRF of the envelope modulations predictor derived from the female speaker stimulus. Notice the lack of a significant response in the average TRF or a region of significance over the null model. Similar results were observed for the carrier stimuli: **Bottom Left:** average TRF of the carrier predictor derived from the male speaker stimulus. Note the significant response in the TRF at the same latency observed for the corresponding envelope TRF. **Bottom Right:** average TRF of the carrier predictor derived from the female speaker stimulus. As in the case of the corresponding envelope TRF, there is no significant response observed for this TRF.

The envelope TRFs for the male speaker exhibited a significant response over the null models driven by an effect from 13 to 43 ms (*t*_max_ = 4.643, *p <* 0.001). Similarly, the significant response of the carrier

TRFs to the male speaker was driven by an effect from 19 to 37 ms (*t*_max_ = 3.393, *p <* 0.001). These results corroborate those obtained in Kulasingham et al. (2020). No significant responses were found for the TRFs for the female speaker.

We used a linear mixed effects model to test the differences between the male and female speaker TRFs. The model was fit to the maximum TRF amplitude for each subject in the range 20-50 ms. The model that best captured the variability in the data (as determined by backward elimination from a maximal model; see section 2.10) was given by:

peak amplitude *∼* speaker gender

i.e., a single fixed effect of speaker gender and no random effects. The effect of speaker gender was significant (*F* = 18.28, *p <* 0.01), indicating that speaker gender was the only meaningful predictor of peak height. Additionally, since the reduced model did not contain any effects of predictor type, we conclude that there is no substantive difference between the envelope modulations and carrier predictors in the single-speaker condition – both contribute significantly to predicting the neural response.

### 3.4 Cocktail-Party TRFs

In the single-speaker case, we reported significant differences in the TRFs of subjects listening to male and female speakers, which differ strongly in their acoustics. In contrast, the cocktail party conditions do not strongly differ in their acoustics but rather only in the subjects’ task and state of selective attention; additionally, TRFs are simultaneously obtained for the male speech and female speech for the same stimulus. We repeated the TRF estimation procedure from the single-speaker analysis, with the result being four average TRFs for the cocktail-party scenario: carrier and envelope TRFs for male attended and unattended speech. TRFs for female speech were estimated but not analyzed further due to lack of a significant response. The grand average TRFs are presented in Figure 4.

**Figure 4.**
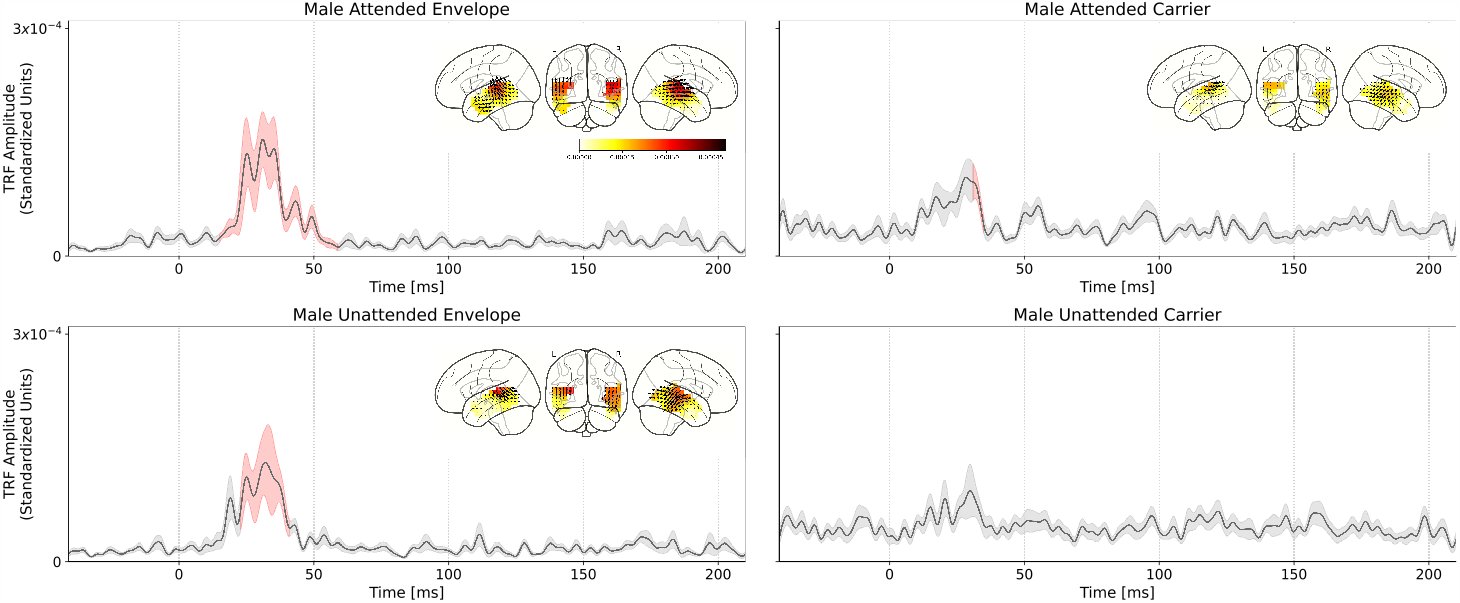
Comparison of attended and unattended TRFs for the male speech stimuli, in the cocktail-party setting. Solid black lines indicate the TRF grand average (over TRF amplitude, averaged across voxels in the ROI); shaded regions indicate values within one standard error of the mean. Red shading indicates TRF values significantly above the noise floor. The distribution of TRF vectors in the brain at the time with the maximum significant response is plotted as an inset for each TRF. **Top Left:** Male speech envelope TRF for subjects attending to the male speech (female speech is background). A large significant response in the TRF is observed between ~30-50 ms which indicates a consistent, time-locked neural response to the speech envelope modulations at a *∼* 30-50 ms latency. **Top Right:** Male speech envelope TRF for subjects attending to the female speech (male speech is background). **Bottom Left:** Male speech carrier TRF for subjects attending to the male speech (female speech is background). **Bottom Right:** Male speech carrier TRF for subjects attending to the female speech (male speech is background). Linear mixed effects model and post-hoc test results indicate that the attended speech TRF peak amplitude is significantly greater than the unattended speech TRF peak amplitude.

As in the single-speaker scenario, we compared TRF amplitudes to those of the average null model to determine the significance of the TRF peaks. Statistical tests revealed that the envelope TRFs for the attend male speaker condition exhibited a significant peak over the null models driven by an effect lasting from 15 to 59 ms (*t*_max_ = 4.230, *p <* 0.001). Similarly, the significant regions of the carrier TRFs to the attend male speaker condition were driven by an effect lasting from 31 to 35 ms (*t*_max_ = 2.755, *p* = 0.04). In the case of the unattended male speaker condition, only the envelope TRF was significant over the null model, driven by an effect lasting from 23 to 31 ms (*t*_max_ = 3.651, *p <* 0.01).

Next, the effect of selective attention on TRF peak amplitude was analyzed. A linear mixed effects model was fit to the maximum TRF amplitude for each subject in the range 20-50 ms. The model that best captured the variability in the data (as determined by backward elimination from a maximal model; see section 2.10) was given by:

peak amplitude *∼* predictor type *×* attention focus + (predictor type | subject)

The model indicates that the fixed effects and interaction of predictor type and the focus of attention (attend male or attend female) significantly contribute to its prediction of the TRF peak amplitudes, even when controlling for variation in predictor response strength at the subject level. A statistical summary for each model is presented in Table 1. The presence of a significant interaction (*t* = *−* 2.499, *p* = 0.012) between predictor type and attention focus suggests that TRF response strength is modulated by attention to different degrees between the envelope modulations and carrier predictors.

**Table 1.**
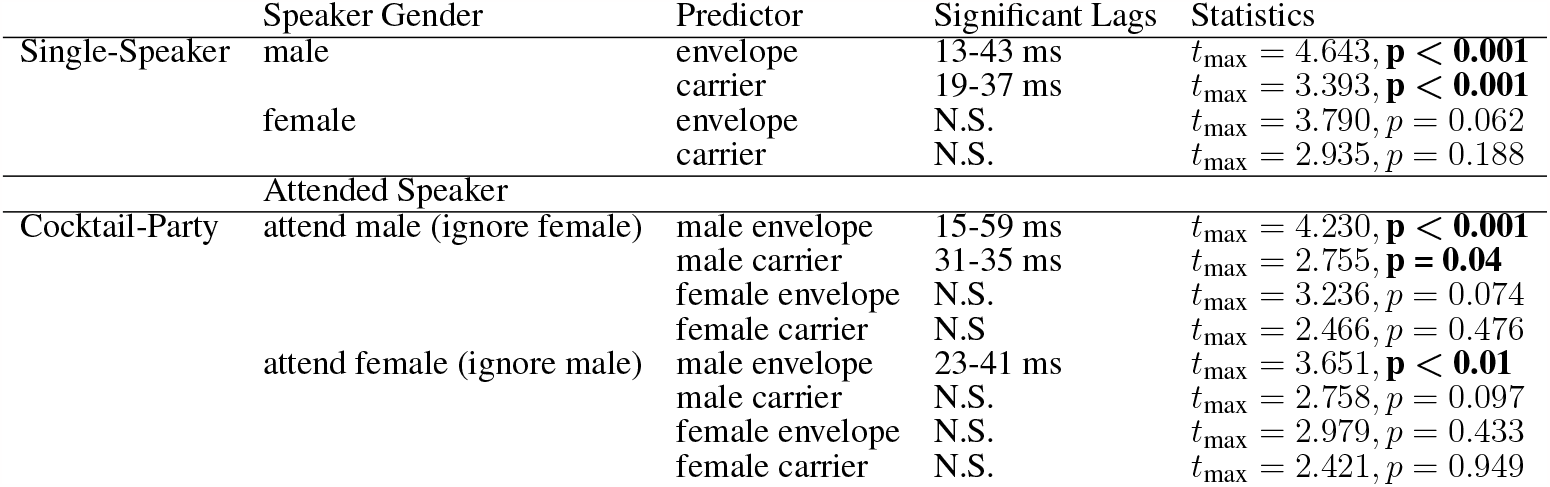
Statistical summary for single-speaker and cocktail-party models.

**Table 2.**
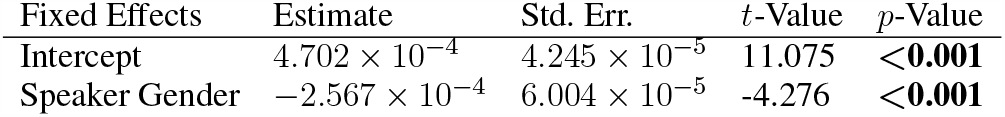
Linear Mixed Effects Model summary, Single-Speaker model.

**Table 3.** Linear Mixed Effects Model summary, Cocktail-Party model.

| Fixed Effects | Estimate | Std. Err. | t-Value | p-Value |
| --- | --- | --- | --- | --- |
| Intercept | $2.159 \times 10^{-4}$ | $4.518 \times 10^{-5}$ | 4.780 | **<0.001** |
| Predictor Type | $1.440 \times 10^{-4}$ | $4.815 \times 10^{-5}$ | 2.990 | **<0.01** |
| Attended Speaker | $2.079 \times 10^{-5}$ | $3.678 \times 10^{-5}$ | 0.565 | 0.57 |
| Predictor Type : Attended Speaker | $-1.300 \times 10^{-4}$ | $5.202 \times 10^{-5}$ | -2.499 | **<0.05** |
| Random Effects | Variance | Std. Dev. |  |  |
| Intercept — Subject | $3.003 \times 10^{-8}$ | $1.733 \times 10^{-4}$ | | |
| Predictor Type — Subject | $2.125 \times 10^{-8}$ | $1.458 \times 10^{-4}$ | | |

A post-hoc Wilcoxon signed-rank test was conducted to test attentional modulation of peak TRF amplitudes between attended and unattended conditions. Two tests were conducted: one for the envelope TRFs and one for the carrier TRFs. The results showed a significant difference for the envelope TRFs (*W* = 29.0, *p <* 0.001) and no significant difference for the carrier (*W* = 122.0, *p* = 0.899). Figure 5 shows individual subjects’ maximum TRF amplitudes in the attend and ignore conditions (male speaker only).

**Figure 5.**
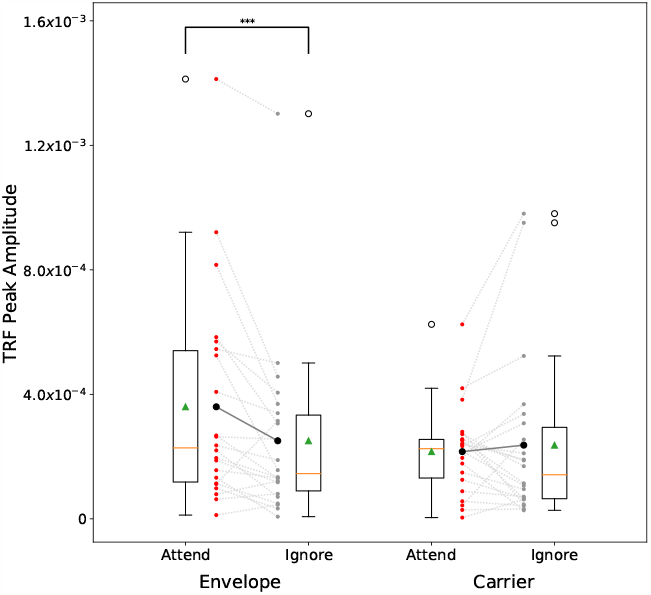
Cocktail-party male speech TRF peak amplitude comparison across subjects. Male speech TRF peak amplitudes in the latency range 20-50 ms are presented for attend male (red) and ignore male (grey) conditions. Dashed lines show each individual subject’s change in peak height between attend and ignore conditions. Solid lines show the change in the mean between the conditions. For the envelope TRFs, note the significant decrease in the mean value, and for most subjects, between the conditions. No such trend is observed in the carrier TRFs.

## 4 DISCUSSION

In this study, we investigated time-locked high-gamma cortical responses to continuous speech measured using MEG in a cocktail-party paradigm consisting of concurrent male and female speech. Such responses were found, and their volume-source localized TRFs provided evidence that these responses are modulated by the focus of attention.

### 4.1 Effect of F0 on High-Gamma Cortical Responses

Most prior studies on high-gamma cortical responses to speech, whether FFR or continuous speech TRFs, employ male speech (e.g., Hertrich et al., 2012; Kulasingham et al., 2020; Canneyt et al., 2021a; Gnanateja et al., 2021; Guo et al., 2021; Kegler et al., 2022; Schüller et al., 2023b). Male speakers typically have lower F0 (;2100 Hz) than typical female speakers with a higher F0 (;2200 Hz). Kulasingham et al. (2020) observed that even for speech stimuli restricted to a single male speaker, the lower pitch segments of voiced speech contributed more to the cortical response than segments with higher pitch. Furthermore, Canneyt et al. (2021b) compared responses to male and female speech, observing that cortical response strength was inversely related to F0 and rate of F0 change throughout continuous speech. Schüller et al. (2023a) recently presented a study wherein gamma-band responses to competing male speakers, with low and high fundamental frequencies respectively, were recorded using MEG. As expected, they reported a significant dropoff in neural response strength to the speaker with the higher F0, a large enough effect that in some cases the responses could not be distinguished from the noise.

In the present study, we have replicated the findings of these previous works by demonstrating a significant difference in the strength of high-gamma cortical responses to male and female speech. As expected, our results show no significant response to female speech, whether in the concurrent speech paradigm or in isolation. In contrast, our findings show a strong, time-locked response to male speech, whether presented in isolation or concurrently with female speech, at a latency of 30-50 ms. This latency is consistent with a neural origin localized to the primary auditory cortex, and when combined with the relative insensitivity of MEG to subcortical sources, bolsters the idea that high-gamma time-locked MEG responses can act as a unique window into primary auditory cortex, without interference from subcortical or other cortical areas (Simon et al., 2022).

Although no significant responses to female speech were observed in our study, this does not imply that such responses are not present. Recent studies have shown that response strength greatly improves for stimuli with strong higher harmonic content. For instance, Guo et al. (2021) recorded strong cortical responses in subjects listening to speech-like harmonic stimuli with a missing fundamental. Canneyt et al. (2021b) also observed that stimuli with strong harmonic content evoke stronger cortical responses.

### 4.2 The Effect of Selective Attention on High-Gamma Cortical Responses to Continuous Speech

In this work, we assessed the effects of selective attention on time-locked high gamma cortical responses to continuous speech. When subjects were instructed to attend to the male speaker, their time-locked responses to the speech envelope modulations and carrier were significantly larger than when subjects ignored the male speaker. As anticipated, no significant time-locked high-gamma responses were seen for the female speaker, either as a single speaker or concurrently with the male speaker. In this way, the use of a female speaker removes any ambiguity as to the source of the neural responses by enforcing a large gap between the competing speakers’ F0 bands. This resulted in enhanced responses to the male speech stream which were then assessed for strength modulation depending on the focus of selective attention.

The MEG-measured TRFs estimated in our study indicate a cortical origin of the responses with a *∼*40 ms peak latency, in line with the findings from earlier studies on time-locked high-gamma auditory cortical responses (Kulasingham et al., 2020; Hertrich et al., 2012) and support the idea that these responses are due to time-locked responses to the fast (*∼* 100 Hz) oscillations prevalent in vowels produced in the continuous speech of a typical male speaker and localized to the primary auditory cortex.

Effects of selective attention on high-gamma EEG FFR have also been observed previously, for non-speech sounds (concurrent amplitude modulated tones) by Holmes et al. (2017), and for simple speech sounds (concurrent vowels) but only when already segregated at the periphery (presented dichotically) (Lehmann and Schönwiesner, 2014). The FFR frequencies for which these selective attention effects were observed (*∼*100 Hz, and 170 Hz, respectively) are consistent with a neural source of primary auditory cortex, but the FFR paradigm does not lend itself to latency analysis. Recently, using TRF analysis of EEG responses to continuous speech, Kegler et al. (2022) demonstrated that a high-gamma TRF (with a latency profile consistent with a contributing source of primary auditory cortex), was modulated by the presence or absence of word-boundaries, i.e., a higher order cognitive (linguistic) cue. Schüller et al. (2023a) also used a TRF approach on MEG data to show that, for male competing speakers, neural responses are modulated by selective attention.

### 4.3 Caveats and Summary

While the speech stimuli were complete sentences and equations, they were not naturalistic continuous speech: the text was spoken at a fixed rate, the sentences were unrelated to each other, and the syntactic/mathematical form of the stimuli was strongly stereotypical. The results seen here would be strengthened by similar findings using continuous speech.

In summary, we have shown that time-locked high-gamma cortical responses to speech are modulated by selective attention in a cocktail-party setting. We have previously argued that time-locked high-gamma MEG cortical responses to speech constitute a valuable physiological window into human primary auditory cortex, with minimal interference from subcortical auditory areas, due to MEG’s relative insensitivity to subcortical structures, and minimal interference from higher order cortical areas, due to the high-frequency/low-latency of the responses. In this way we provide new evidence for (top-down) selective attentional processing of competing speakers as early as primary auditory cortex.

